# A defensive body plan was pre-adaptive for termitophily in the rove beetle tribe Termitohospitini (Staphylinidae: Aleocharinae)

**DOI:** 10.1101/083881

**Authors:** T. Kanao, K.T. Eldredge, M. Maruyama

## Abstract

Termitophily—the symbiosis of organisms with termite societies—has evolved a disproportionate number of times within the rove beetle subfamily Aleocharinae (Staphylinidae). Among aleocharine termitophiles, defensive (limuloid) and mimetic (physogastric & physothoracic) body forms have evolved convergently, but due to lack of a comprehensive aleocharine phylogeny, the context in which termitophily and associated adaptations evolve is unknown. We present the first example of a robust, morphology-based phylogenetic placement of an exclusively termitophilous tribe, the Termitohospitini. Termitohospitini is recovered to be nested within Myllaenini *sensu nov*, and sister to Myllaena (new synonymy). Furthermore, we also recovered the small tribe Masuriini nested within Myllaenini *sensu nov* (new status).

Reconstructing ecological transitions within this clade, we present evidence that the stem lineage of Myllaenini sensu nov invaded intertidal marine habitats, the common ancestor for Myllaena + Termitohospitini then transitioned to freshwater riparian habitats, with Termitohospitini alone subsequently shifting to termitophily. We conclude that: (1) Termitohospitini was ancestrally a limuloid-bodied riparian inhabitant; (2) a limuloid form may have been pre-adaptive for defense against host attack during the evolution of termitophily; (3) the strongly tapered abdomen of an ancestral limuloid body was a constraint on the evolution of physogastry, leading to the emergence of the unusual physothoracic body form observed in termitohospitines that likely integrates these obligate termitophiles to life inside termite colonies.

“one of the most astonishing spectacles in all natural history.” — Richard Dawkins

## Introduction

Some of the most ecologically complex and taxonomically diverse organismal interactions center around the societies of social insects, in particular those of ants and termites (Rettenmeyer et al. 2011, Kistner 1979, 1982, Hölldobler & Wilson 1990, Parker 2016). Organisms that have evolved symbioses with ants and termites are termed myrmecophiles and termitophiles, respectively, and as a group these symbionts display a bewildering array of morphological and behavioral specializations. Among termitophiles, two body forms, mimetic (Figs. 8–9B) and defensive (Figs. 1–7), have convergently evolved numerous times. Mimetic species typically possess swollen “physogastric” or “physothoracic” (Figs. 8–9B) body forms— the consequence of increased fat body growth that results in a grotesquely expanded body, stretching out the intersegmental membrane between sclerites. Physogastry is thought to transform the sclerotized body into one that resembles the relatively soft-bodied host termites (Cunha et al. 2015), often to a striking degree (Dawkins 1996, Kistner 1969). In such species, body segments begin to distend soon after the adults eclose (post-imaginal growth) (Kistner 1969), and in some instances elongation of leg segments even occurs, breaking the cardinal rule that arthropods grow by moulting. In defensive, or limuloid (horseshoe crab shaped) groups, the abdomen is posteriorly tapered, lateral margins of the body are typically expanded and head deflexed, protecting vulnerable appendages dorsally from attack (Figs. 1–4, 6–7) (Kistner 1979, Parker 2016).

**Figures 1–8.**
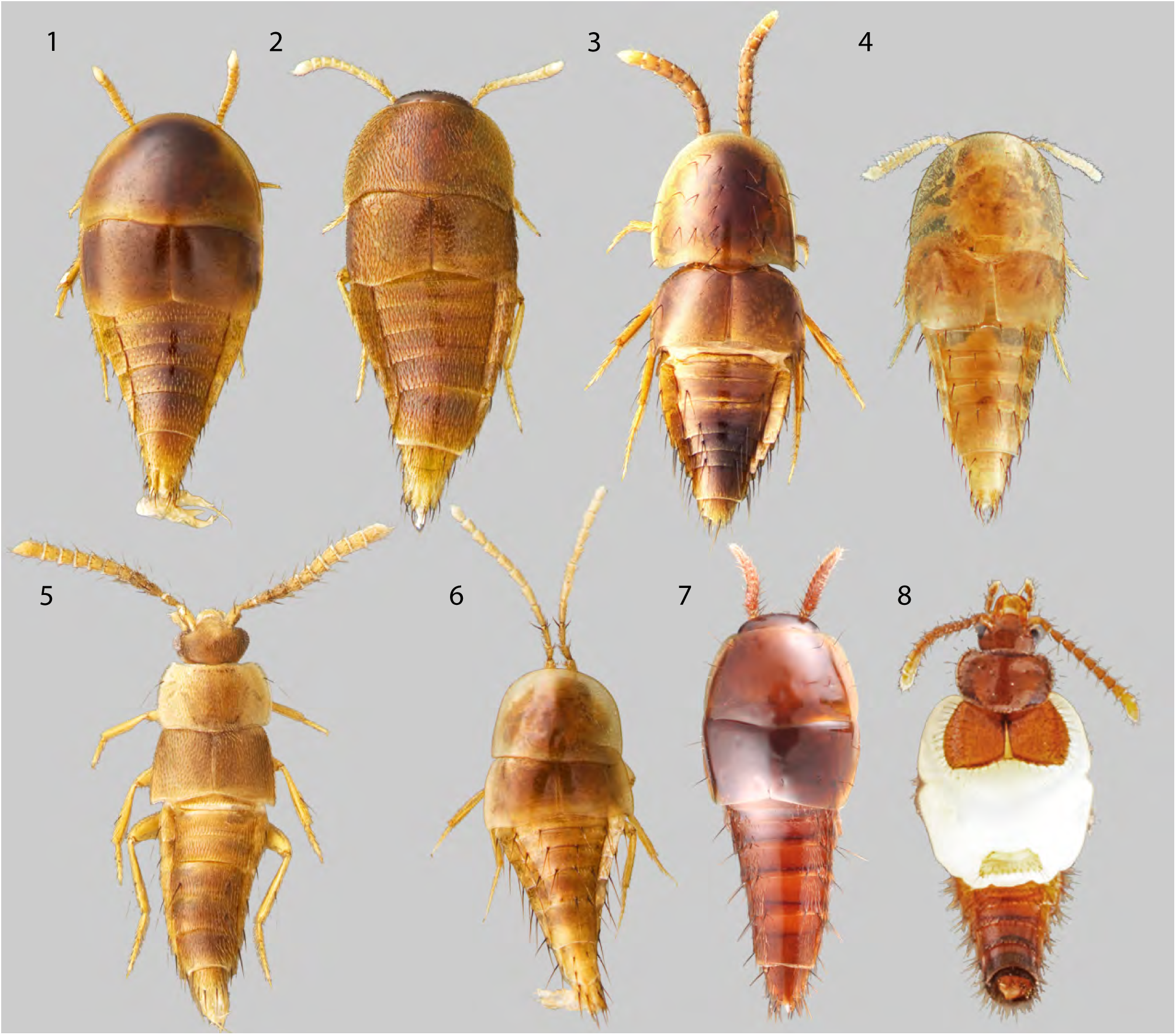
Habitus of representative Termitohospitini. (1) *Termitohospes* undesc. sp. (2) *Paratermitosocius* undesc. sp. (3) *Termitosodalis* undesc. sp. (4) *Termitosocius microps*. (5) *Neotermitosocius bolivianus.* (6) *Blapticoxenus brunneus*. (7) *Coptotermocola clavicornis*. (8) *Coptoxenus* sp.

**Figures 9–10.**
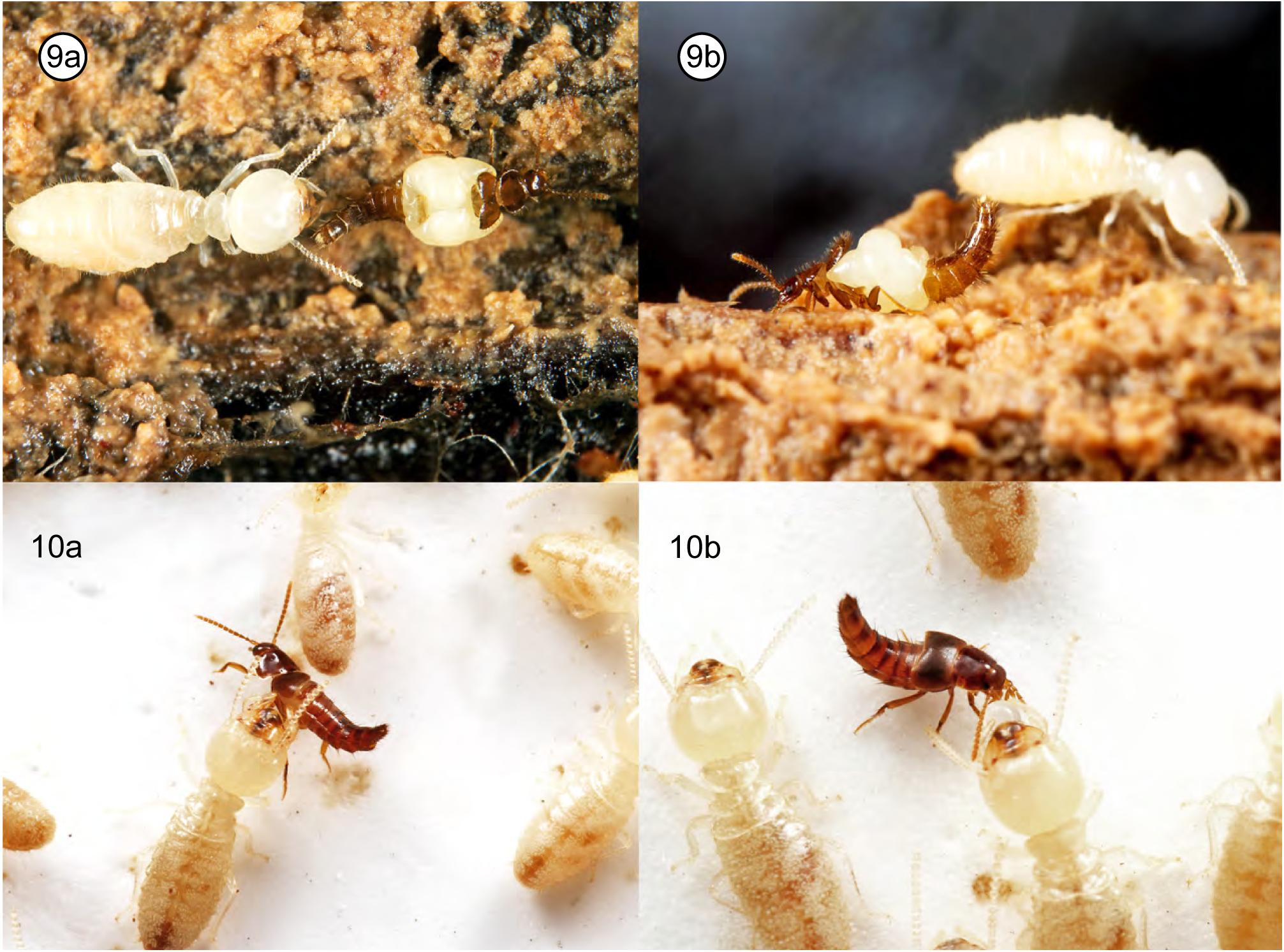
Termitohospitini in situ et in vitro. (9) *Coptoxenus* sp. in situ with host. (10a) *Sinophilus* sp. in vitro being groomed by host. (10b) *Sinophilus* sp. in vitro engaging in trophalaxis food exchange.

Across the Arthropoda, termitophilly has evolved only rarely, if at all (REF ####). One exception is the rove beetle subfamily Aleocharinae, where this lifestyle has arisen at least 11 times, and over 650 described species are known to be symbiotic with termites (Kistner 1969, Kanao et al. 2012). Such an exceptional predisposition to evolving this type of symbiosis raises an immediate question over potential traits that might be pre-adaptive for the aleocharine capacity to successfully invade termite nests. One approach to identifying putative pre-adaptations is to explore the evolutionary interplay between morphological and ecological specializations leading up to independent origins of termitophily across the Aleocharinae phylogeny. However, a major impediment exists in the lack of phylogenetic resolution in Aleocharinae, and in the social insect symbiont lineages in particular (Ashe 2007). Consequently, we have little understanding of the ecological contexts within which termitophily evolves among aleocharines, and the character transformations leading to the specialized morphologies seen in many termitophilous clades. This phylogenetic impediment is particularly due to the taxonomic challenges one confronts with Aleocharinae; their small size often obscure species level differences, and vast taxonomic diversity impedes their study (there are 16,468 described species, making them the largest animal subfamily [Newton unpublished data]) (Ashe 2007). Compounding the situation, social insect symbionts are notoriously challenging to collect due to their low abundances (similar to army ant symbionts [Kistner 1979]) and difficulty in extracting them from inside nests (Kistner 1998, personal observations).

During a survey of aleocharine morphology, conducted for a comprehensive morphological treatment of the subfamily (Eldredge in prep.), we discovered that the lacinia (a food gathering structure of the mouthparts [Betz et al. 2003]) of one specialized termitophilous tribe, Termitohospitini (Figs. 1–10B, S4), is strikingly similar to Myllaenini (Figs. 11A–11B, S4), a tribe notable for genera that have specialized on marine-intertidal and freshwater-margin habitats, but which are entirely free-living. Furthermore, a previous study has suggested that the particular lacinial morphology of Myllaenini is in part characteristic for the tribe (Ahn & Ashe 2004). Despite the marked ecological disparity between termitohospitines and myllaenines, further examination revealed the genus Myllaena (Myllaenini) and many termitohospitines shared a unique sensory organ on the aboral surface of maxillary palpomere 3 (Figs. 12A–12B). Therefore, we hypothesized a priori that a phylogenetic analysis would reveal the inclusion of Termitohospitini within a broader Myllaenini, and a sister-group relationship between Myllaena + Termitohospitini. Here, we present the first instance of a robust phylogenetic hypothesis concerning the common ancestry of a termitophilous aleocharine lineage, Termitohospitini, providing a unique opportunity to explore the sequence of morphological and ecological character evolution leading to the origin of symbiosis in a termitophilous aleocharine lineage.

**Figures 11–13.**
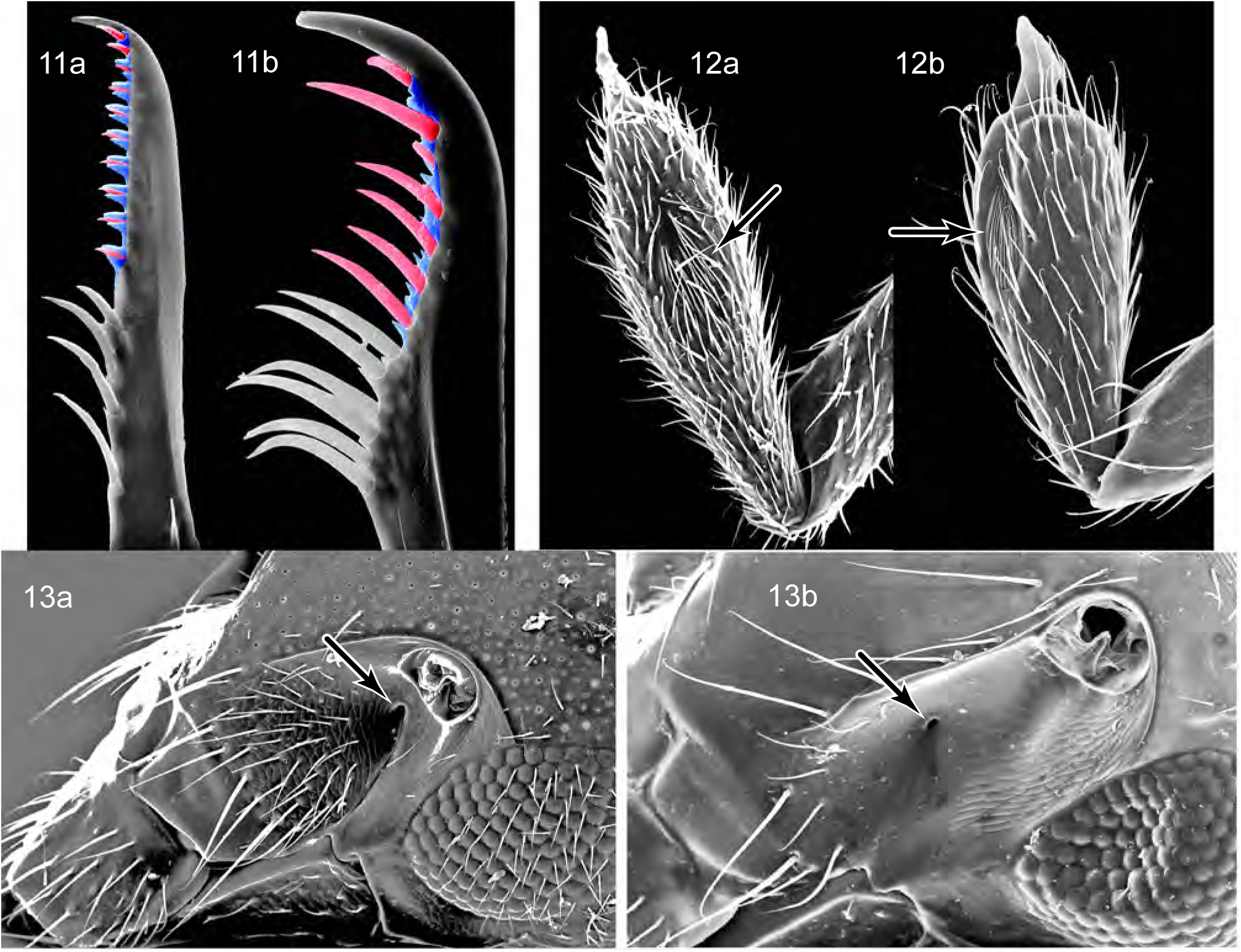
*Myllaena* sp. and Termitohosptini scanning electron micrograph images of various body parts. (11a) *Myllaena* sp. acinia. (11b) *Sinophilus* sp. lacinia. (12a) *Myllaena* sp. maxillary palpomere 3. (12b) *Termitohospes* sp. maxillary palpomere 3. (13a) *Myllaena* sp. anterior tentorial pit. (13b) *Termitohospes* sp. anterior tentorial pit.

## Materials and Methods

### Specimen Work

Specimens for this study were made available by the Snow Entomological Museum Collection (University of Kansas) and numerous field expeditions conducted by Kanao and Maruyama (also see Acknowledgements). We again emphasize that symbionts of social insects are exceedingly rare and the difficult to collect. Symbionts number one per thousands of colony members, and nests are often subterranean and inaccessible (Kistner 1979, 1998).

To examine the morphology, typical procedures were taken of KOH tissue digestion, chlorazol black staining, examination in glycerin, and permanent mounting in Euparal (for details see Eldredge 2012 Hanley & Ashe 2003, Maruyama 2004). Light- (Olympus SZX7 stereomicroscope and Olympus BX51 compound microscope) and scanning electron microscopy was used for morphological examination. Whole body images were taken using a Visionary Digital™ BK Plus Lab System with Cannon EOS 7D. Images were captured from multiple focal planes and combined with CombineZM (http://hadleyweb.pwp.blueyonder.co.uk/CZM/News.htm).

### Dataset Construction

For this study, we modified the dataset of Ahn & Ashe (2004). Sixteen taxa and nine additional characters were added for a total matrix dimension of 74 taxa by 108 characters (S1–2). The dataset was constructed and handled in Mesquite v3.1.0 (Maddison & Maddison 2016).

### Phylogenetic Analyses

Parsimony analyses were done with TNT v1.1 (Goloboff et al. 2000) implementing ratchet and drift algorithms to increase exploration of tree space, and 1,000 bootstrap replicates were performed. Analyses were done locally.

The Bayesian reconstructions were analyzed in MrBayes v3.2.6 (Ronquist & Hulsenbeck 2003) using the Mkv model (Lewis 2001). Several strategies to account for among site character rate differences were implemented by modifying the Mkv model. Gamma rates with four and eight rate categories were implemented in MrBayes v3.2.6 hosted by CIPRES Science Gateway (Miller et al. 2010), and a log-normal distribution was implemented with a modified version of MrBayes (Harrison & Larsson 2015) hosted by KINBRE (University of Kansas). Analyses were run for at least 50 million generations and was monitored with Tracer v1.5 (Rambaut & Drummond 2007) to confirm ample mixing. Sampled trees were summarized with DendroPy v3.12 (Sukumaran & Holder 2010).

PAUP* v4.0b10 (Swofford 2002) and WinClada v1.00.08 (Nixon 2002) were used to plot apomorphies. Ancestral state reconstructions were performed using a Mk likelihood reconstruction available in Mesquite v3.1.0.

We initially began by reanalyzing the Ahn & Ashe (2004) dataset with the objective of adding termitohospitine representatives for preliminary insight. During this course, we discovered errors in the published dataset, preventing reconstruction of the presented results. Through author correspondence the source of error was identified and corrections are reflected in the dataset presented in this study (S2).

## Results

### Phylogeny Reconstruction

#### Parsimony

106 most parsimonious trees of 785 steps were recovered (majority rule consensus tree, Fig. 14). Termitohospitini was recovered monophyletic with strong support (92% bootstrap), notably supported by the anterior migration of the anterior tentorial pits (char. 99:1) and antennae set in acetabullae (char. 100:1) (Figs. 13a–13b, S6–7). Further, Termitohospitini was recovered sister to Myllaena, which in turn is sister to Dimonomera and nested deep within the Myllaenini. The sister-group relationship of Termitohospitini and Myllaena was found to be supported by the presence of a unique sensory structure on maxillary palpomere 3 (char. 103:1) (Figs. 12a–12b).

**Figure 14.**
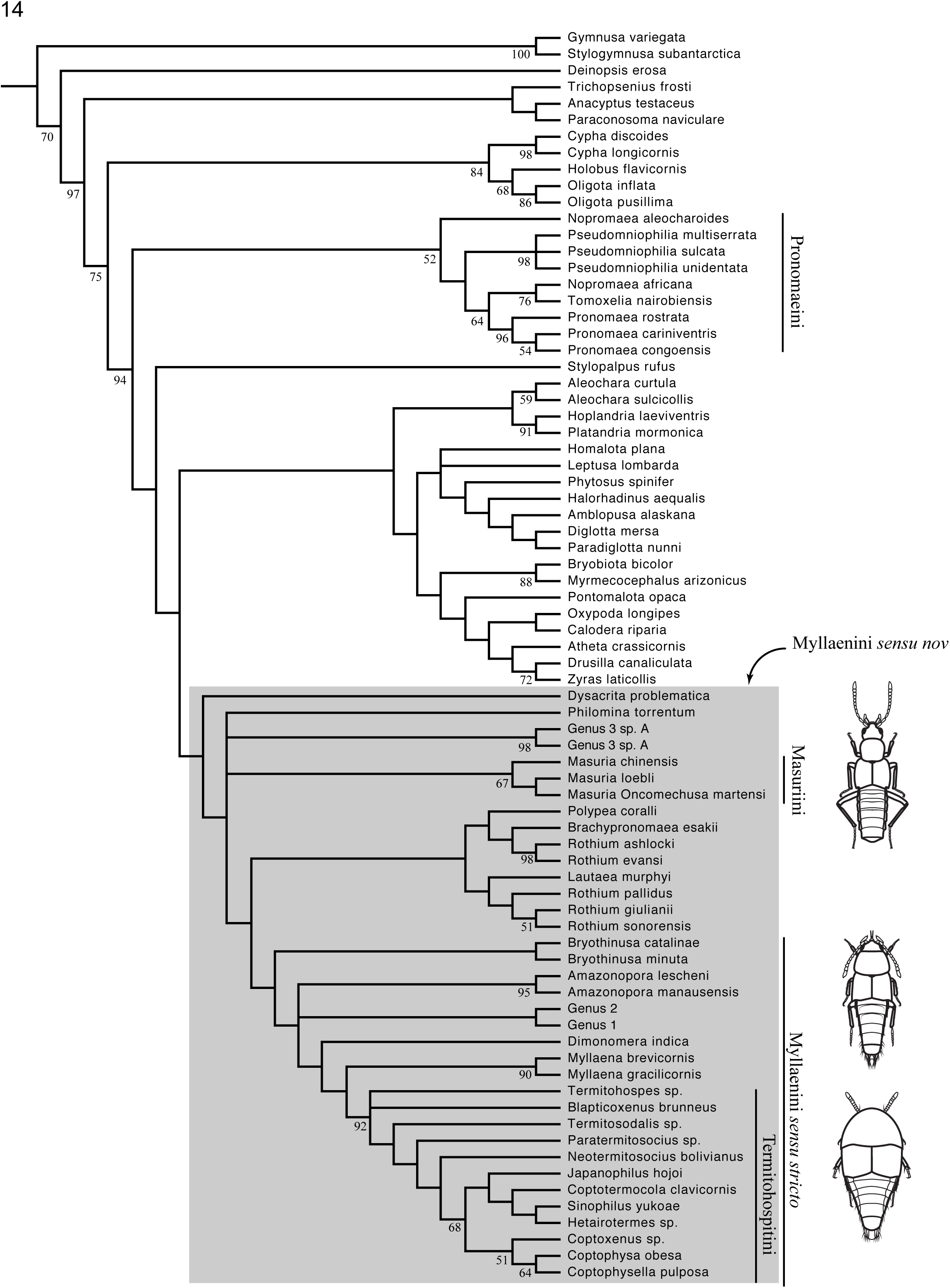
Parsimony majority rule consensus tree with bootstrap values.

A clade consisting of Myllaenini sensu Ahn & Ashe was recovered to share a most recent common ancestor with Dysacryta and Masuria (Masuriini): Myllaenini sensu nov. Myllaenini sensu nov is supported by, among others, the unique lacinial morphology of large interdigitating setae (char. 26:2) (Figs. 11A–11B, S4). Pronomaeini was not recovered sister to Myllaenini sensu nov.

The genus Rothium was rendered paraphyletic by Lautaea murphyi, which formed a clade with R. pallidus, R. giulianii and R. sonorensis; Brachypronomaea esakii and Polypea coralli formed a clade with R. ashlocki and R. evansi. The Lautaea clade is not supported by a clear synapomorphy, but the Brachypronomaea clade is supported by the mentum being medially squarely produced (char. 108:1).

A clade consisting of Bryothinusa, Genus1, Genus2, Amazonopora, Dimonomera, Myllaena, and Termitohospitini was recovered in all analyses: Myllaenini sensu stricto, which is supported by a four-articled mid tarsi (char. 105:1) among other characters, but no unambiguous changes support this node.

The monophyly of Bryothinusa was unresolved in the Bayesian analyses, but was recovered in the parsimony analysis. Bryothinusa monophyly is supported by an incomplete infraorbital carina (char. 5:1) and quadrate pronotum (char. 57:3).

Very little support was recovered overall. Most of the recovered nodes failed to generate any support, and those that did were concentrated at the deepest nodes and tips.

#### Bayesian

The general results were similar to those recovered from a parsimony analysis (Figs. 15–16). Termitohospitini was recovered monophyletic and sharing a MRCA with Myllaena and Dimonomera in all analyses with strong support. The general topology of Myllaenini sensu stricto is similar with the parsimony results, but less resolved. In particular, the Bayesian analyses failed to support the monophyly of Bryothinusa which remains unresolved. In all the analyses, Lautaea was recovered sister to Genus 3, differing from the parsimony analysis.

**Figure 15.**
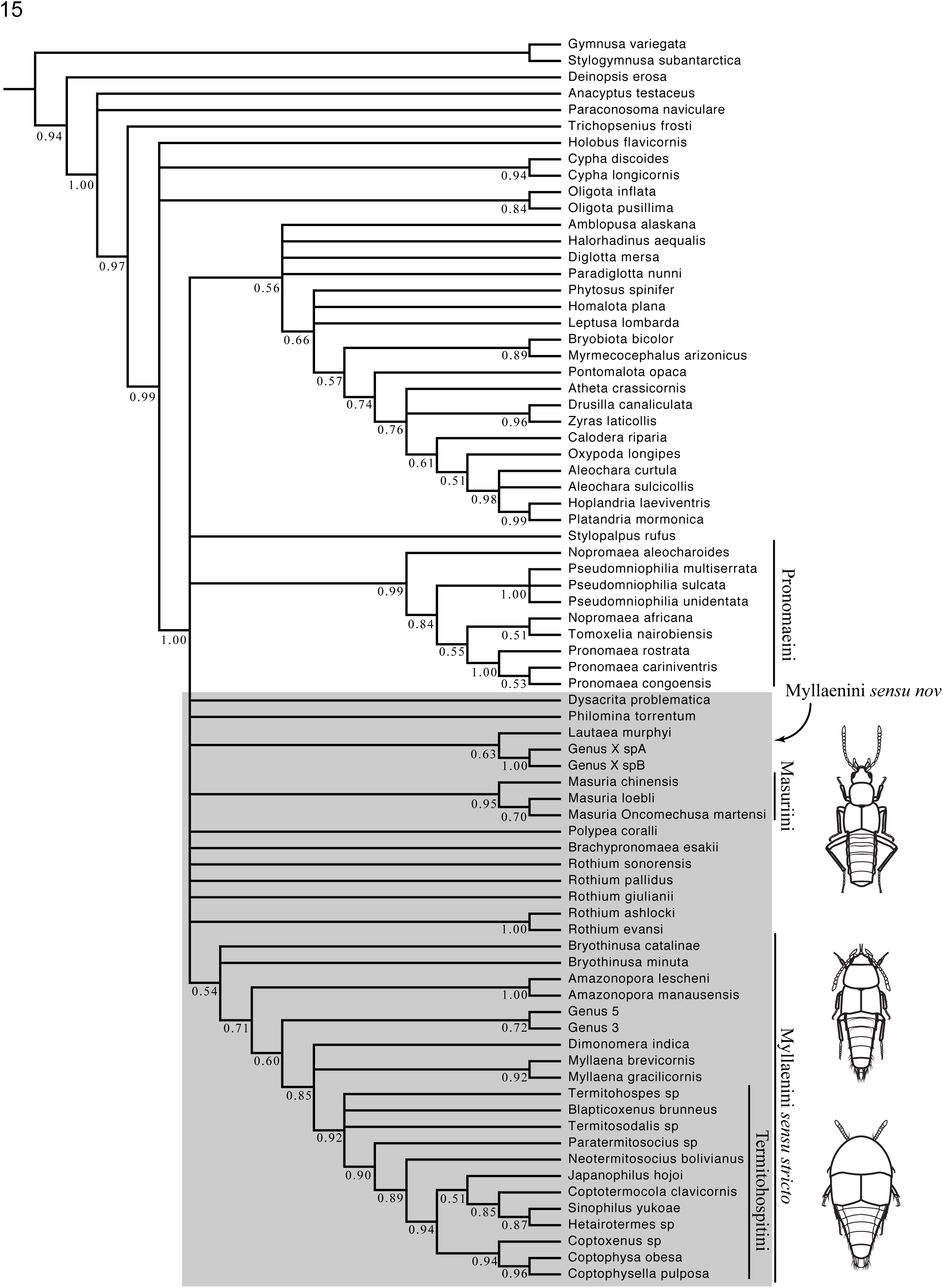
Bayesian Mkv + Γ(4) majority rule consensus tree with posterior probabilities. Habitus illustrations from top to bottom: *Masuria* sp., *Myllaena* sp. and *Termitohospes* sp.

**Figure 16.**
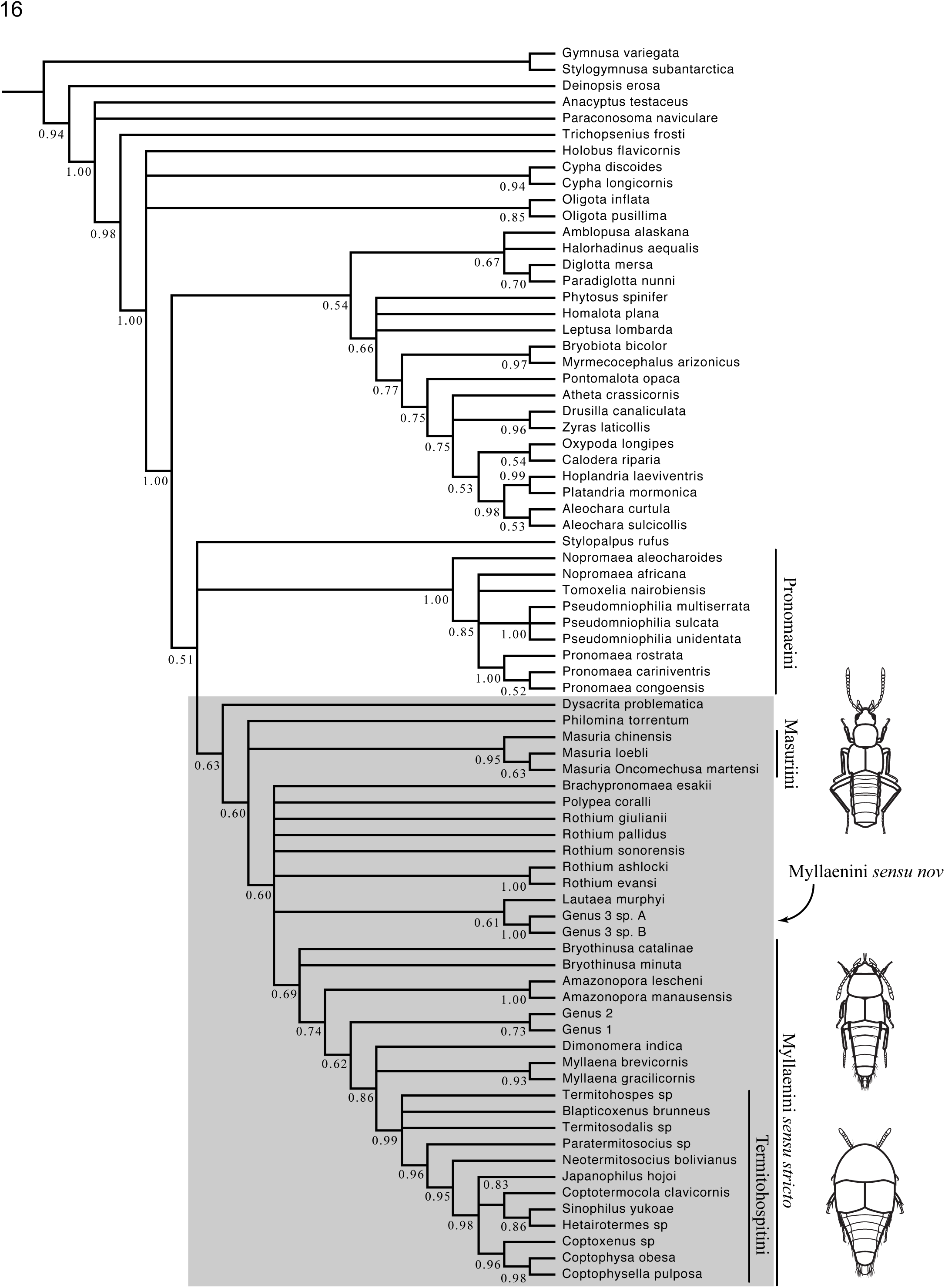
Bayesian Mkv + Γ(8) majority rule consensus tree with posterior probabilities. Habitus illustrations from top to bottom: *Masuria* sp., *Myllaena* sp. and *Termitohospes* sp.

**Figure 17.**
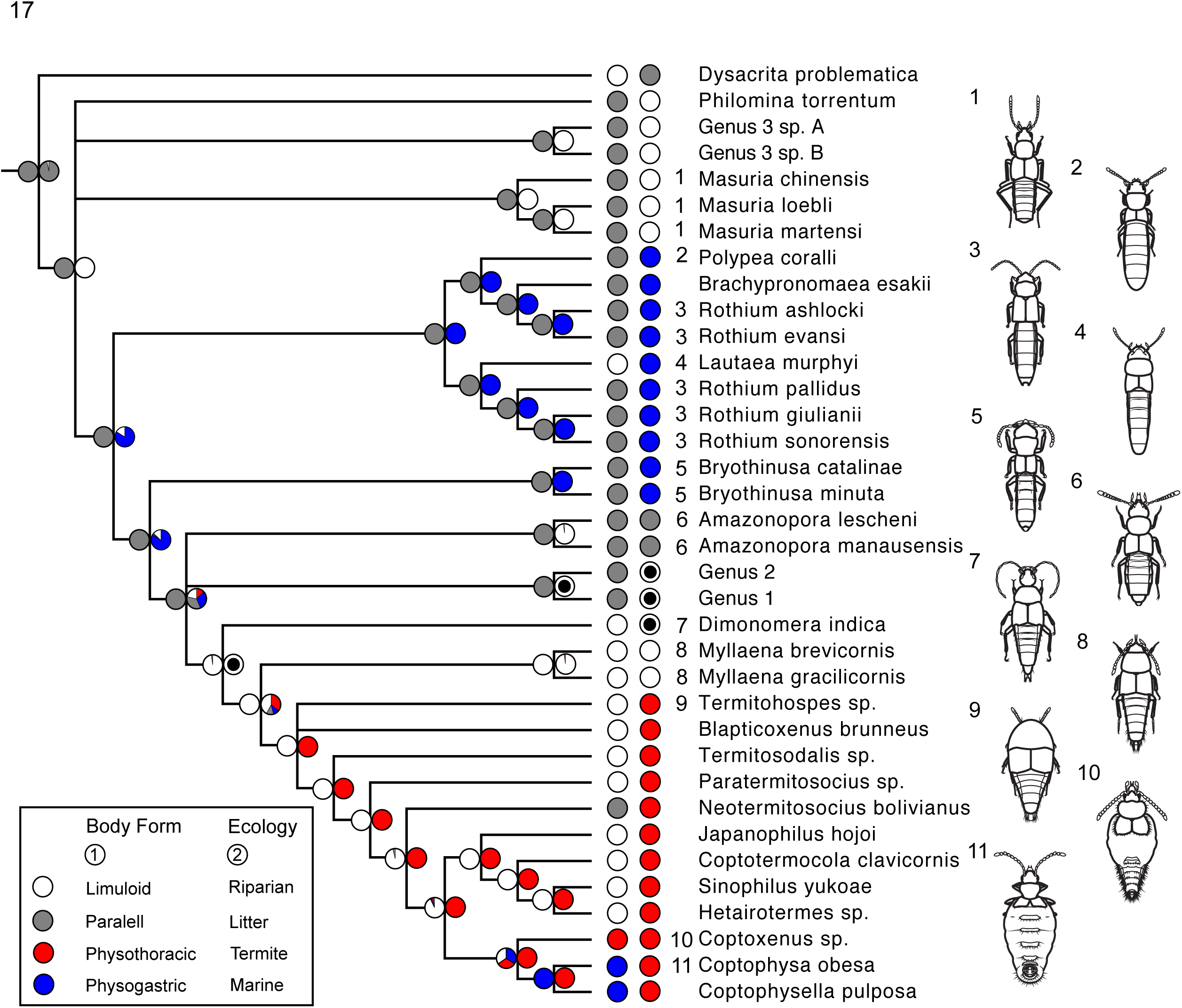
Parsimony majority rule consensus tree of Myllaenini *sensu nov* with body form and ecology Mk reconstructions.

In the Mkv+G4 analysis (Fig. 15), asides from Rothium ashlocki + R. evansi, Masuria and Lautaea + Genus3, a significant proportion of Myllaenini sensu nov members were unresolved in their placement.

In the Mkv+G8 analysis (Fig. 16), Myllaenini sensu nov was recovered but with little support. Asides from Rothium ashlocki + R. evansi, the placement of Rothium, Polypea and Brachypronomaea was unresolved.

### Character Evolution

Ground/leaf litter habitat was recovered ancestral for Myllaenini sensu nov (Fig. 18). The group subsequently transitioned from riparian to marine, and reverted back to riparian habitat before infiltrating termite nests with Termithospitini. Myllaenini sensu nov is ancestrally parallel-sided. Limuloid form evolved twice, once in the mangrove swamp inhabiting Lautaea murphyi and the common ancestor of Dimonomera + Myllaena + Termitohospitini, transitioning to physogastry and physothoracy within the termitohospitines. The ancestral state for the MRCA of Coptoxenus + Coptophysa + Coptophysella is ambiguous.

### Taxonomy

Taxonomy proposed by Ahn & Ashe (2004):

Myllaenini sensu Ahn & Ashe
  = Dimonomerini (Ashe 1999)
    Amazonopora
    Brachypronomaea
    Bryothinusa
    Dimonomera
    Dysacrita
    Lautaea
    Myllaena
    Philomina
    Polypea
    Rothium

The following updated taxonomic scheme is proposed following the results of this study:

Myllaenini *sensu nov*
  = Dimonomerini (Ashe 1999)
  = Masuriini (new status)
  = Termitohospitini (new synonymy)
    *Amazonopora*
    *Brachypronomaea*
    *Bryothinusa*
    *Dimonomera*
    *Dysacrita*
    *Lautaea*
    *Masuria*
    *Myllaena*
    *Philomina*
    *Polypea*
    *Rothium*
    *Blapticoxenus*
  Termitohospitina
    *Coptophysa*
    *Coptophysella*
    *Coptotermocola*
    *Coptoxenus*
    *Hetairotermes*
    *Japanophilus*
    *Neotermitosocius*
    *Paratermitosocius*
    *Sinophilus*
    *Termitobra*
    *Termitohospes*
    *Termitosodalis*

## Discussion

### Ecological Origins of Termitophily

Termites are generally ground inhabiting social insects. By extension of the ground habitat, some inhabit dead wood, but the majority build subterranean nests that may be accompanied by above-ground architecture. Therefore, we logically expected that termitophilous lineages would have a similar ancestral ecology, living on the ground/leaf litter or in dead and decaying wood. Counter to this notion, we recovered a freshwater-riparian habitat-type for the common ancestor of Myllaena + Termitohospitini. This is surprising, since riparian habitats are notoriously underutilized by termites (as well as by ants). A fraction of Myllaena inhabit ground/leaf litter in the tropics, and it is possible that these ground environments are quite saturated, acting as transitional habitats between riparian zones and more strict terrestriality. We hypothesize that termitohospitines evolved from ancestors that ventured into soaked ground/leaf litter habitats in a tropical environmental setting. Furthermore, we posit that certain extant Myllaena lineages are currently experimenting with this riparian-to-ground habitat transition.

### Pre-adaptatation and Subsequent Constraint of the Limuloid Body Plan

Although we observe a parallel-sided body as ancestral for Myllaenini sensu nov, reconstruction shows that Termitohospitini evolved from a limuloid ancestor. Dimonomera and Myllaena are very similar in form overall, which suggests that Termitohospitini evolved from a Myllaena-like, limuloid-bodied ancestor. For riparian limuloid aleocharines like Myllaena, the limuloid body plan is used for pushing, wedging and weaving through substrate. Analogous examples are Gymnusa and Deinopsis, early diverging (Ashe 2005) free-living riparian aleocharines that have similar body forms and live in similar habitats. There appears to be a negative correlation between “limuloid-ness” and abdominal flexibility. The more tapered the abdomen, the less flexible it becomes. Again, Gymnusa and Deinopsis represent analogues of this phenomenon.

In contrast, limuloid bodies in social insect symbionts are adaptive for defense against host attack. The numerous examples of symbiotic aleocharines converging on limuloid ecomorphology are strong evidence of its adaptive nature (Parker 2016). In many examples, the dorsum of the body is broadly expanded laterally, shielding vulnerable appendages underneath. Termitohospitini were ancestrally limuloid but the mimetic physogastric or physothoracic shape, evolved from this limuloid ancestral condition. It is likely that the ancestral limuloid body plan was pre-adaptive for termitophily in the termitohospitines, catalyzing diversification in this niche. Such a scenario parallels a recent hypothesis for the evolution of termitophily elsewhere in Aleocharinae: in the tribes Trichopseniini and Mesoporini, quasi-limuloid morphology of a Gymnusa-Deinopsis-like ancestor is posited to have been preadaptive for specialization inside termite colonies (Yamamoto et al. 2016).

Interestingly, termitohospitines represent the sole example of physothoracy in the Aleocharinae (Figs. 8–9B). There may be several explanations for this uniqueness. Physothoracy may be a strategy to achieve mimicry when the abdomen is too inflexible to easily evolve physogastry. Indeed, a relatively inflexible abdomen appears to be correlated with the limuloid body plan that termitohospitines inherited from a Myllaena-like ancestor. Alternatively, physothoracy may represent an intermediate state toward full physogastry, or perhaps functions as a wholly different function inside colonies, unrelated to social integration. Due to the limited number of species of mimetic termitohospitines, it is presently difficult to establish the polarity of this character, which might help distinguish between these alternatives. Consequently, reconstruction of the MRCA of the mimetic termitohospitine clade remains unresolved (Fig. 18).

### Mouthpart Degradation: A Result of Adaptation to Trophallaxis

A conspicuous characteristic of Myllaenini sensu nov is their extremely stylate mouthparts that greatly extend past the apex of their head (i.e. clypeus). Although all of the mouthpart components are to some extent elongated, the most noticeable are the maxilla and labium. Observations of Myllaena in captivity revealed that the labium, which protrudes past all other mouthparts, is a highly extrusible structure. At rest, the labium is typically retracted into the recesses of the oral cavity, and does not visibly protrude, which is made possible by the elongate premental tendons (Eldredge pers. obs.).

The maxilla is the primary appendage used in food gathering, and the labium is primarily used for sensing. Observations of Bryothinusa have shown these beetles are carnivorous and also scavenge on small arthropods between sand particles in marine-intertidal habitats (Wong & Chan 1977). Similarly elongate mouthparts are convergently found in the riparian limuloid genera Gymnusa and Deinopsis. Observations of Gymnusa showed these beetles are carnivorous, and when feeding, probed the food item with the labium and gathered food with the maxilla (Eldredge pers. obs.). Using these examples of correlated form and function as evidence for a dietary type, Myllaenini appears to be a predaceous tribe; the flexible and elongate labium is probably used to probe substrate interspaces to search for food, and the stylate maxilla used for extracting interstitial food items.

The large interdigitated teeth are a synapmorphy for Myllaenini sensu nov (S5), but many of the termitohospitines have much shorter and curved maxillae (Figs. 11a–11b, S5).

Degradation of the mouthparts is a common theme among social insect symbionts, and is thought to be correlated to a simplified diet: feeding on soft-bodied brood or directly fed by workers.Until recently, the diet of Termitohospitini was unknown, but observations of beetles in culture have revealed that Termitohospitini are fed trophallactically by host termite workers, and also groom and are groomed by the workers (Figs. 10a–10b). Grooming host termites may also allow the beetles to collect colony-specific odors. We suggest that degradation of mouthpart morphology is adaptive for this transition from a predatory diet to one based around trophallaxis and grooming. Notably, the Termitohospitini genus Termitosocius possesses the elongate, more stylate mouthparts characteristic of free-living Myllaenini sensu nov. We therefore speculate that Termitosocius is early-diverging among Termitohospitini, retaining the ancestral predatory diet within termite nests.

### Putting It All Together: Evolutionary Origins of Termitophily in Termitohospitini

Our reconstructions show that termitohospitines evolved from a riparian limuloid ancestor with stylate mouthparts. We hypothesize that transitioning to a more terrestrial environment caused an increase in the frequency of interactions with termites, and the limuloid defensive form was pre-adaptive for protection against termite aggression. Termitohospitini further modified the ancestral limuloid form by laterally expanding dorsal sclerites to shield vulnerable appendages underneath (Figs. 1–4, 6–7). Subsequently, some lineages evolved trophallactic behavior Fig. 10b), and the mouthparts degraded and shortened in length. Given this hypothesis, Termitosocius appears most primitive in overall morphology, and may embody what a stem termitohospitine might have looked like during the lineage’s ecological transition to life inside termite colonies.

## ACKNOWLEDGEMENTS

Collectively we thank Dr. Kee-Jeong Ahn for graciously providing us with original dataset, Dr. Zack Falin and Dr. Floyd Shockely for access to institutional collections, Dr. Takahashi Komatsu for image use, Mr. Taku Shimada for image use and sharing valuable observational data, and Dr. Joseph Parker for critical review of a draft.

